# Expanding the size limit of RNA viruses: Evidence of a novel divergent nidovirus in California sea hare, with a ~35.9 kb virus genome

**DOI:** 10.1101/307678

**Authors:** Humberto J Debat

## Abstract

While RNA viruses thrive with massive structural and functional diversity, their genomes size variation is particularly low, ranging only from ~2-to-33 kb. Here, I present the characterization of RNA sequences corresponding to the first virus associated with *Aplysia californica*. Genome structure and domain architecture suggest that the identified virus is a novel member of *Nidovirales*. The proposed aplysia californica nido-like virus (AcNV), with a genome sequence of ca.35,906 nt, represents the longest ever recorded RNA virus yet. Phylogenetic insights indicate that AcNV clusters in a major phylloclade of unclassified invertebrate nidoviruses, *Roniviridae*, and *Mesoniviridae*. Basal branching in this emerging cluster could indicate that AcNV is a member of a novel divergent clade within *Nidovirales*. Further, virus RNA detection in multiple independent studies suggests that AcNV is neurotropic with a broad cell/tissue/organ tropism, supported by AcNV occurrence in diverse organs, including the first detection of a *Nidovirales* in single specific neurons.

**Highlights:**

-RNA virus genomes reported in the literature are limited at ca. 33.4 kb
-A novel nidovirus was identified in the gastropod mollusk *Aplysia californica*
-The aplysia californica nido-like virus (AcNV) presents a 35.9 kb RNA genome
-AcNV has a broad tropism, is enriched in the CNS, and accumulates in neurons
-The unique features of *A. californica* enables single-neuron virus dynamics of AcNV

## 1. Introduction

A massive flood of RNA sequence data is becoming accessible as a result of the exponential expansion and democratization of high-throughput sequencing platforms. Metatranscriptomics studies are exploring and elucidating the expression profiles of a global plethora of organisms. In tandem, a growing multifaceted RNA virosphere is emerging, uncovering the viral dark matter associated with every single living organism assessed (Greninger, 2018). Despite the exponential growth of data generation during the last years and the parallel discovery of thousands of new emergent viruses, one crucial characteristic of RNA virus genomes, which has maintained constant in the metagenomics revolution (perhaps the only constant), is an upper-limit on genome length of <34 kb in RNA viruses (Shi et al., 2018). Ball python nidovirus (*Nidovirales, Coronaviridae*) presents a 33.4 kb RNA virus genome (Stenglein et al., 2014) which still remains to be the largest reported to date and has been associated with respiratory disease in experimental infections of ball pythons (Bodewes et al., 2014; Hoon-Hanks et al, 2018). Nidoviruses are positive-stranded RNA enveloped viruses with a monosegmented large genome. The linear nidoviruses genome is infectious, and its RNA molecule is capped and polyadenylated. The order *Nidovirales* is comprises four different families: *Arteiivindae, Mesoniviridae, Coronaviridae* and *Roniviridae* (Adams et al., 2017). While genome size of nidoviruses differs significantly amid ca 12.7-33.4 kb, a general conserved genome organization, their replication strategy and sequence similarity in the replicase protein suggest that nidoviruses share a common ancestor (Gorbalenya et al., 2006). Nidoviruses are widespread in humans and numerous animal species, and have been detected in terrestrial and marine mammals, fish, birds, reptiles, insects and crustaceans (De Groot et al., 2012; Nga et al., 2011; Lauber et al., 2012). Interestingly, despite their evident broad host range, only one nidovirus has been associated with mollusks (Phylum Mollusca), more specifically with a *Turritella* sea snail (*Gastropoda*) (Shi et al., 2016). The California sea hare (*Aplysia californica*) is a sea slug marine opisthobranch gastropod mollusk in the family *Aplysiidae*. This gigantic gastropod (up to 75 cm and 7 kg of weight) and valuable laboratory animal presents a “simple” nervous system of about 20 thousand neurons, distributed in 10 ganglia of ca. two thousand cells each. These neurons are of massive size and present differential pigmentation, which has been useful for specific individual identification. Targeted studies with individual neurons of *A. californica* had led to the understanding of specific contributions of concerted neurons to simple and complex behaviors in which they are involved. Hallmarks of behavior neurochemistry have been elucidated in this gastropod such as the classic siphon-withdrawal reflex documented by Eric Kandel’s fundamental early works, demonstrating for the first time that environmental cues may generate structural effects on the nervous system (Carew et al., 1972; Kandel, 1976). Here, I present the genomic characterization of the first virus to be associated with *A. californica*, a tentative novel member of order *Nidovirales*, with the largest ever-recorded RNA virus genome.

## 2. Results

### 2.1 Virus discovery in A. californica

In order to assess the potential RNA virus landscape of *A. californica*, every available RNA resource at NCBI was employed for virus discovery. 117 RNA NGS libraries, of diverse studies, target organ/tissue/cell, origin or developmental stage, encompassing over 164.54 gigabases were explored. The raw reads were downloaded and subjected to *de novo* assembly with Trinity v2.6.6 release or the CLC genomics workbench v8.0.3 assembler into library specific transcriptomes which were further subjected to bulk BLASTX searches (E-value < 1e-5) against a local database of refseq virus proteins. The obtained hits were explored by hand. Interestingly, a large 35.9 kb transcript assembled from a single-cell RNAseq library generated from the interneuron L-29 (SRA = SRR3211539) got a significant best hit (E-value = 7e-22; 24 % identity) to the orf1ab gene product (YP_005352837) of *White-eye coronavirus HKU16*; (Nidovirales; Coronaviridae; *Deltacoronavirus*). Iterative mapping of the interneuron L-29 reads were employed to polish the identified transcript which was curated into 35,906 nt, derived of 2,235,790 RNA reads (5.86 % of total library reads) giving an average coverage of 6382,4X. Surprisingly, equivalent or shorter contigs sharing ≥99 % identity to the detected transcript were found in 61 additional *A. californica* generated transcriptomes. Moreover, raw read mapping of all available libraries using as cut-off >10 reads with a ≥99 % identity suggests that this RNA is present in 71 independent NGS libraries, which is roughly 60.6 % of all available *A. californica* RNA libraries. Additional analyses were carried out to rule out that this transcript could be derived from the expression of an endogenous *A. californica* gene (see section 2.3).

### 2.2 Structural and functional annotation of Aplisya californica nido-like virus (AcNV)

Further structural characterization and annotation of the detected transcript indicate the presence of two major ORFs (Figure 1.A). ORF1a (67-17,521 nt coordinates) appears to be followed by a putative pseudoknot (Figure 1.E) that could be associated to translation read-through of a UGA stop codon (opal) at position 17,519-21 nt to generate a larger ORF1ab by suppression of termination (67-25,198 nt coordinates). After a 53 nt spacer a second large ORF2 is predicted at 25,252-34,920 nt coordinates. The encoded ORF1ab and ORF2 are flanked by a 67 nt A/U rich (61.2 %) 5’UTR and a 964 nt long 3’UTR followed by a 22 nt Poly (A) tail. This genome organization, while clearly different from the hallmark configuration of nidoviruses, is shared with the only available nido-like virus associated with a gastropod reported yet: Beihai nido-like virus 1 (BNLV-1; Shi et al., 2016). BNLV-1 presents two large ORFs encoding two polyproteins, and the first polyprotein encoded in ORF1ab is tentatively generated by read-through of a premature stop codon at 7708-7710 nt, instead of a characteristic −1 ribsomal frameshifting, the standard for nidoviruses (Brierley et al., 1987). In the *A. californica* detected RNA sequence, the ORF1ab encodes a polyprotein (PP1ab) of 8,375 aa. To my knowledge, besides the recently described Diaphorina citri flavi-like virus and Gamboa mosquito virus polyproteins (8,960 aa and 8,572 aa, respectively) this is the third largest ever-predicted protein for any RNA virus yet. Blastp searches of PP1ab against the non-redundant protein sequences (nr) obtained as best hit the orf1ab polyprotein of *Bulbul coronavirus HKU11-796* (E-value = 5e-36; 23 % identity). ORF2 (25,252-34,920 nt coordinates) encodes a highly divergent polyprotein (PP2), with no evident virus best hits, but a meager similarity (E-value = 0.035) to the spike glycoprotein of *Ball phyton nidovirus 1*. Further structural and functional *in silico*annotation of the PP1ab and PP2 proteins evidenced the presence of several domains in a canonical position and a general architecture typical of nidoviruses (Figure 1.A.*i*). Three major hydrophobic regions comprising multiple trans membrane-spanning regions were predicted with TMHMM v2.0 in ORF1a PP (TM1 _1607_L-Y_1820_, TM2 _4174_F-F_4286_, and TM3 _4704_T-Y_4884_) (Figure 1.A.*iiiii*). A putative membrane protease like region (PL, proteolysis associated domain; YdiL; E-value = 1.60e-03) was identified among 1603-1760aa positions. Further, between TM2-TM3 a 3C-like proteinase (E.C.3.4.22.-) region was spotted by HHPred (3CL; PDB structure: 3D23_A of *Human coronavirus HKU1* main protease; Probability: 96; E-value = 0.045) at 4,293-4,699 aa coordinates. This main protease of nidoviruses, flanked at both sides by TM2-TM3, similar to that of picornavirus 3C proteinases, is responsible for the processing of the most conserved part of the replicase polyprotein (ORF1b) (Anand et al., 2002). In the ORF1b polyprotein region, HHPred searches identified at 6496-6911 aa coordinates of ORF1ab a RNA-directed RNA polymerase (RdRP, PDB structure: 1RAJ_A; Probability: 96.82; E-value = 9.6e-6; Pfam = pfam00680) and at 7,259-7596 aa coordinates a Middle East respiratory syndrome coronavirus helicase domain (HEL, PDB structure: 5WWP_A; Probability: 99.76; E-value = 6.4e-23). This peculiar position of the RdRP upstream the HEL domain is distinctive, exclusive of nidovirales among ssRNA(+) viruses (Gorbalenya et al., 1989), and has also been reported in tentative ssRNA(+) hypovirus-like mycoviruses (Marzano et al., 2016). Additionally, between the RdRP and HEL domains, as expected, a multinuclear coronavirus zinc-binding domain (CV-ZBD) was identified (6,963-7,050 aa; score = 10.893; Prosite = PS51653). The predicted CV-ZBD domain contains 10 His/Cys typical residues at equilocal conserved positions of several nidoviruses (**Supp. Fig 2**). The ZBD domain has been indicated as a genetic marker of order *Nidovirales*, since it has not been identified in any other viral orders (Gorbalenya et al., 1989). The RdRP-ZBD-HEL predicted region of the ORF1ab protein appears to be the more similar to reported nidoviruses (Fig 1.D). Lastly, a divergent methyltransferasa region, similar to that of nsp14 or nsp10, exoribonuclease and transferase (MT; PDB structure: 5C8T_B of Human SARS coronavirus guanine-N7 methyltransferase; Probability: 90.11; E-value = 0.12) was found at the 7,642-7,937 aa coordinates. In sum, the predicted ORF1ab protein presents an array of characteristic domains in an architecture typically associated to nidoviruses (NH2-PL-TM1-TM2-3CLpro-TM3-RdRP-ZBD-HEL-MT-COOH) this specific domain order is essential for diverse aspects of the replicative cycle of nidoviruses (Gorbalenya, 2001). Further characterization of ORF2 encoded protein (PP2) suggests a clear divergence between this polyprotein and ordinary nidoviruses. Nevertheless, divergent but consistent signatures were identified which supports its affinity with BNLV-1. ORF2 encodes a 3,224 polyprotein with an N-terminal signal peptide with a predicted cleavage site between residues Ala_21_ and Gly_22_ (Fig.1.A.*i-ii*). HHPred searches identified at 276-471 aa coordinates of PP2 a Trypsin (TryP; E.C.3.4.21.4) serine protease domain (PDB structure: 5KWM_A; Probability: 98.88; E-value = 4.3e-12). At an equilocal position (128-308 aa coordinates), the ORF2 product of BNLV-1 presents a related domain (Trypsin-like serine protease; Superfamily = SSF50494; E-value = 5.9e-07). While proteases involved in post-translational processing of PP1ab are a hallmark of nidoviruses, evolutionary insights of the detected domain suggest that this peptidase is significantly related to the ones found in the digestive system of many vertebrates, where they hydrolyzes diverse proteins. Future studies should explore the potential source of the detected TryP domain, which might be associated to horizontal transmission from an eventual host. In addition, the C-region of ORF2 of BNLV-1 presents an alphavirus glycoprotein like domain (Superfamily = SSF56983; 1,335-1,546 aa). The *A. californica* ORF2 product presents an analogous, but highly divergent region with similarity with Torovirinae spike glycoprotein (Sgp; 2,185-3,092 aa; FFAS03 = PF17072.3; 17 % id) at the C-region of the predicted PP2, which could be associated with structural functions.

**Figure 1.**
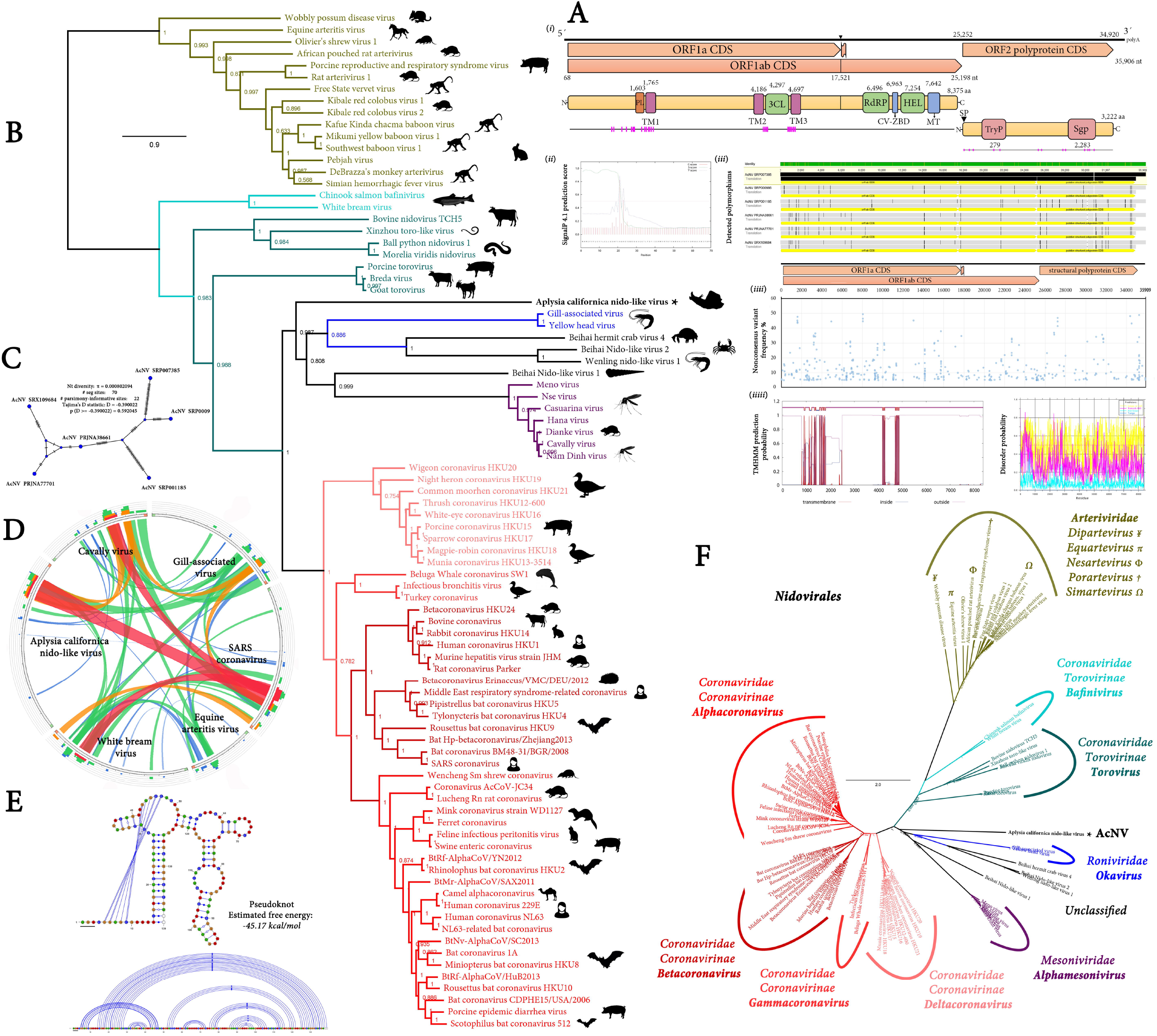
Structural characterization and phylogenetic insights of aplysia californica nido-like virus (AcNV) **A**) (*i*) Genome graphs depicting architecture and predicted gene products of AcNV. Predicted domains are shown in initialized curved rectangles and indicated at their starting coordinate. Flag signal indicates a putative pseudoknot (**E**) that could be associated to translation read-through of a UGA stop codon (opal, black rectangle) at position 17,519-21 nt to generate the ORF1ab polyprotein by suppression of termination. Pink bars indicate transmembrane predicted regions. Abbreviations: TM1-3, trans-membrane domains 1-3; 3CL, 3C-like proteinase (3CL^pro^); CV-ZBD, multinuclear coronavirus zinc-binding domain; PL, proteolysis associated domain; RdRP, RNA-dependent RNA polymerase; HEL, helicase domain; MT, methyltransferase-like domain; SP, signal peptide cleavage site; TryP, Trypsinlike serine protease domain. (*ii*) SignalP-4.1 prediction of the cleavage site (Ala_21_/Gly_22_) of ORF2 polyprotein. C-score, raw cleavage site score; S-score, signal peptide score; Y-score, geometric average of the C-score and the slope of the S-score (*iii*) Detected polymorphic sites in consensus sequences of diverse *A. californica* bioprojects generated with a >75% frequency call per-base. Variable sites are indicated by black lines in both the genomic RNA sequence track (top) and/or the predicted gene products (below) of each consensus. (*iiii*) Non-consensus variant frequency as a percentage of total mapped bases per site determined with 625,367,351 base calls of 5,796,298 AcNV virus derived RNA reads. Detailed polymorphisms are available in **Supp. Table 4**. (*iiiii*) Multiple membrane-spanning regions of ORF1ab replicase determined with TMHMM v2.0 (left). Intrinsic disorder regions (right) predicted with the disEMBL tool. Pink lines, Remark-465 predictor; turquoise, hot-loops; yellow, loops or coils. **B**) Maximum likelihood phylogenetic tree based on MAFFT alignments of predicted replicase protein of AcNV (black star) and related *Nidovirales*. The *Arteiiviridae* family was used as tree root. Genera of depicted viruses are indicated by colors following (**F**). Scale bar represents substitutions per site. Node labels are FastTree support values. Silhouettes indicate the reported host of the respective nidovirus. **C**) Tentative haplotype network based on PopART using the integer neighbor-joining algorithm. Each blue vertex represents a tentative viral haplotype. Branch bars represent the number of mutations between sequences. **D**) Similarity levels of AcNV and representative members of each *Nidovirales* family or sub-family replicases expressed as Circoletto diagrams based on BLASTP searches with an E-value of 1e-1 threshold. RPs are depicted clockwise, and sequence similarity is visualized from blue to red ribbons representing low-to-high sequence identity. **F**) Unrooted radial phylogenetic tree visualization of (**B**), genera of depicted viruses are indicated by colors.

Overall, genetic similarity, genomic organization, functional and structural annotation suggest that the identified transcript could be sequence evidence of a new species of RNA virus with cues of *Nidovirales*. Thus, I propose the name Aplisya californica nido-like virus (AcNV) to the detected putative virus.

### 2.3 Genomic evidence supports AcNV as bona fide extant virus associated to A. californica

Using as query the predicted gene products of AcNV, multiple TBLASTN searches (E-value >1e-1) targeting the *A. californica* genome assembly AplCal3.0, NCBI accession number GCF_000002075.1 failed to return any significant hits. This result supports the proposition that the AcNV sequence does not derive from the RNA expression of an integrated DNA element sharing similarities with *Nidovirales*, or contamination of RNA sequencing libraries with *A. californica* DNA harboring such element, or potential AcNV like endogenous viral elements EVEs (Katzourakis & Gifford, 2010). Further, the complete raw DNA reads used to generate the *A. californica* genome were explored in detail (Bioproject PRJNA13635), to rule out that the AcNV sequence could correspond to an unassembled region of the *A. californica* genome. In this direction, MEGABLAST of the raw DNA reads (E-value >1e-1) with AcNV resulted in no significant hits to the virus sequence. In sum, these data are in concordance with the proposal that the detected AcNV RNA sequence is, in fact, evidence of a *bona fide* extant virus associated to *A. californica*.

### 2.4 Phylogenetic insights of AcNV

To entertain the hypothesis that the proposed AcNV is related to the *Nidovirales* order, phylogenetic insights were generated based on MAFFT multiple amino acid alignments of predicted ORF1ab proteins of accepted and proposed members of *Nidovirales* (**Supp. Table 1**), followed by FastTree maximum likelihood phylogenetic trees (Fig 1.B, F). Several major lineages could be identified in the obtained phylogenetic tree of *Nidovirales*: (*i*) A subfamily *Coronavirinae* (*Coronaviridae*) lineage which includes exclusively vertebrate viruses, and can be divided in 4 genera (*Alpha-Betta-Gamma-Delta-coronavirus*). (*ii*) A lineage of small nidoviruses (*Arteviridae*) associated also to vertebrates, which has been recently separated into five genera (*Dip-Equ-Nes-Por-Sim-arterivirus*). (*iii*) A lineage of fish (*Bafinivirus*) and snake and mammals viruses (*Torovirus*) which are included in a diverse *Coronaviridae* subfamily (*Torovirinae*). (*iiii*) Lastly, a cluster of medium sized nidoviruses, enriched in mosquitoes (*Mesoniviridae*) and a crustacean group of viruses (*Roniviridae*) in a concerted lineage with divergent crustacean and a mollusk virus which remain unclassified, could be identified. The obtained tree indicates that AcNV unequivocally clusters within the order, in the pivotal major clade composed of recently proposed crustacean and gastropod nidoviruses, the *Roniviridae*family and *Mesoniviridae*. The local topology of the tree and basal branching of AcNV could suggest that this *A. californica* associated virus might be the first member of a new lineage of nidoviruses (Fig. 1.B).

### 2.5 AcNV predicted prevalence, tropism and variability

AcNV could be traced to expression sequence tag (EST) libraries generated more than 10 years ago (Moroz et al., 2006; Fiedler et al., 2010). Over 97 ESTs were further linked to AcNV, presenting highly supported similarity to the virus sequence (E-value ≤ 9e10-72; identity ≥ 92%). The tentatively AcNV derived ESTs were generated from libraries obtained from the central nervous system (CNS) of *A. californica*, most hits being linked to a specific pedal and pleural ganglia sample library (**Supp. Table 2**). The suggestive hints towards a CNS enrichment inclined me to further study the potential tropism of AcNV.

In light of the unique *A. californica* biology of well-established model organism, its amazing central nervous system, and the availability of vast genomic resources, I explored the genomic RNA landscape of AcNV in Californica sea hare. The distinguishing neuroanatomy of *A. californica* and its well-studied neurons offer an exceptional technical platform for RNA dynamics (Puthanveettil et al., 2013; Moroz & Kohn, 2010–2013). To tentatively assess cell/tissue and organ tropism based on virus-derived RNA accumulation in *A. californica*, available California sea hare RNAseq libraries were mapped to AcNV and normalized as Fragments Per Kilobase of transcript per Million mapped reads (FPKM). Virus RNA transcripts were detected in an outstanding diversity of target tissue and organs, suggesting e pervasive tissue tropism of AcNV (Fig 2; **Supp. Table 3**). At the adult organ level (Fig 2.A, B right panel) AcNV was detected in chemoreceptive areas (rhinophores & tentacles), salivary glands, gills, heart, additional muscles (body wall, buccal and penial), hermaphroditic glands, ovotestis, hepatopancreas and other digestive glands. Interestingly, AcNV appeared to be highly enriched in the CNS. The availability of CNS tissue and single-cell RNAseq data allowed the surveillance of specific tropism at the cellular level of AcNV (Fig 2.B-E). AcNV accumulated at relatively higher levels in the left excitatory interneuron 29 (L29), the left motor neuron 7 (L7), the right serotonergic cerebral ganglion metacerebral neuron (MCC-R) and the right abdominal ganglion cholinergic neuron 2 (R2). The R2 neuron and its axonal pathway is highly conserved and thus can be consistently identified in closely related species of the genus *Aplysia*. R2 measures up to 1.1 mm in diameter and is the largest neuron in the animal kingdom (Moroz & Kohn, 2013). AcNV was found at lower relative levels in the left abdominal ganglion motor neuron (LFS), the sensory neuron cluster and the left MCC.

**Figure 2.**
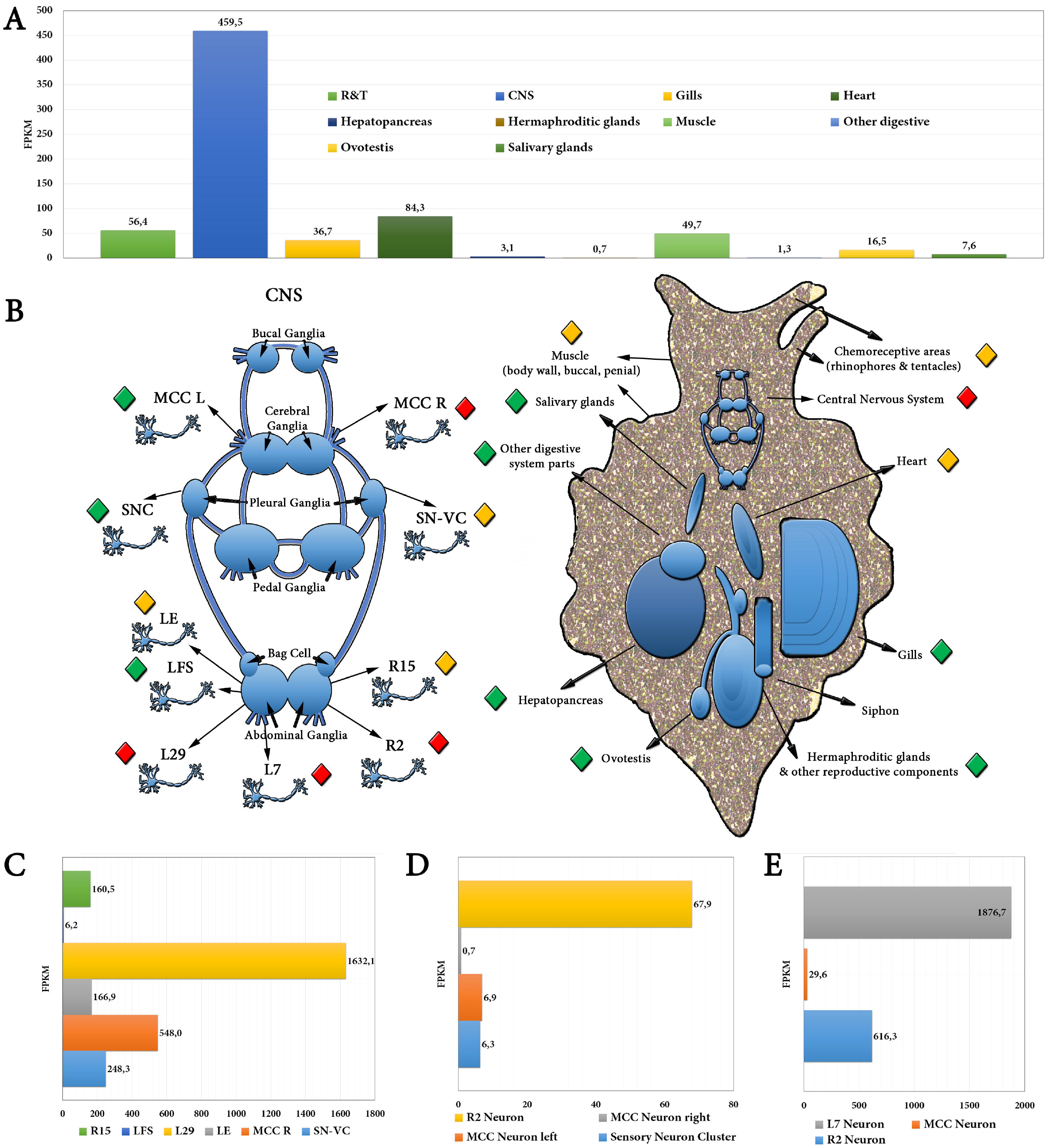
AcNV tentative cell, tissue and organ tropism based on virus-derived RNA accumulation in *A. californica*. **A**) Graphs bars expressing virus RNA transcript levels assayed in diverse expression profile libraries derived from distinct *A. californica* tissues or organs. Values are expressed as Fragments Per Kilobase of transcript per Million mapped reads (FPKM). **B**) *A. californica* diagram (right) highlighting the virus RNA positive tissue/organs and a schematic diagram of the central ganglionic nervous system of *A. californica* (left) indicating specific neurons which presented AcNV virus RNA. RNA levels are visualized as diamonds colored green-yellow-red representing low-to-high relative RNA virus accumulation. Abbreviations: CNS, central nervous system; SNC, sensory neuron cluster; R2-R15, right abdominal ganglion cholinergic 2 and 15 neurons; L7, left motor neuron 7; MCC, serotonergic cerebral ganglion metacerebral neuron; LFS, left abdominal ganglion motor neuron; LE, left excitatory abdominal sensory neuron; L29, left excitatory interneuron 29; SNV-VC, pleural ventrocaudal sensory neuron. C) Graphs bars expressing virus RNA transcript levels assayed in diverse single-cell expression profile libraries derived from specific *A. californica* individual neurons. Virus FPKM values corresponding to each *A. californica* sample are available as **Supp. Table 3**.

In addition to adult organ tissues and cells, I was able to detect AcNV in libraries generated from individuals at very early developmental stages (Heyland et al., 2011) such as early cleavage (4-7 h after egg masses are laid) and third day individuals (1-4 cells; transition from trochophore to veliger body plan). AcNV virus FPKM values corresponding to each *A. californica* sample are available as **Supp. Table 3**.

The vast collection of NGS data allowed both the generation of consensus sequences for each bioproject, based on >75 % base calls, and the identification of variable sites at each position (Fig 1.A.*iiii*). The analyses of every NGS library allowed insights into the variation within-host and eventual evolutionary hints at a genome wide level. The predicted AcNV *bona fide* variation was noticeably spotted across the genome with specific regions, such as the 3’ UTR showing greater levels of nonconsensus calls (**Supp. Table 4**). In order to explore evolutionary relationships among potentially distinct virus lineage populations derived from diverse *A. californica* studies harboring AcNV, haplotype networks were inferred based on multiple MAFFT alignments of complete consensus AcNV virus sequences derived from *A. californica*Bioprojects employing the PopART (Population Analysis with Reticulate Trees) software. The haplotype networks suggest that the variability among consensus sequences of the diverse AcNV virus positive studies is non-significant (Fig. 1.C; Fig.1.A*iii*).

## 3. Discussion

### 3.1 AcNV as a divergent nido-like virus

The AcNV genomic plan, predicted gene products, canonical array of conserved domain of nidoviruses in the ORF1ab protein (NH2-PL-TM1-TM2-3CLpro-TM3-RdRP-ZBD-HEL-MT-COOH), and overall sequence similarity with reported nidoviruses, were considered to propose that the detected RNA sequence corresponded to a nido-like *bona fide* virus. Further, phylogenetic insights indicated that AcNV unequivocally clustered within the order *Nidovirales*, in the pivotal major clade composed of recently proposed crustacean and gastropod nidoviruses, the *Roniviridae* family and *Mesoniviridae*. The basal branching of AcNV could suggest that this *A. californica* associated virus might be the first member of a new lineage of nidoviruses. The discovery of new related viruses is fundamental to unravel the emergent viral dark matter that would license a better understanding of the evolutionary relations and concomitant taxonomic assignment of AcNV in the context of *Nidovirales*. Given the evident association that could be traced in terms of evolutionary links between *Nidovirales* and their hosts, perhaps an interesting follow up strategy would be the (overlooked) exploration of mollusks as source of viral diversity.

### 3.2 RNA virus genome size constrains, AcNV and Nidovirales

AcNV presents the largest RNA virus genome ever recorded (**Supp. Fig 1.A-C**). At ca. 35.9 kb, AcNV is over 2.4 kb longer that the second largest: Ball phyton nidovirus 1 (BPNV-1) RNA genome (Stenglein et al., 2014). When comparing the 84 refseq genomes of nidovirus available at NCBI and their taxonomic assignment to specific *Nidovirales* families or sub-families, three major groups became evident: (i) *Arteriviridae* members, “small nidoviruses”; (ii) *Mesoniviridae*, “intermediate nidoviruses”, and (iii) *Coronaviridae* and *Ronaviridae* “large nidoviruses” (**Supp. Fig 1.A-B**). Within the *Coronaviridae* group, particular dynamics in size may be observed such as a wider pattern of longitudes of subfamily *Torovirinae* members presenting a either ca. 20 kb genomes or larger ca. 32-33 kb ones, the latter being the snake-hosted BPNV-1 and Morelia viridis nidovirus. On the other hand, sub-family *Coronavirinae*members present a more restricted pattern of genome length, which may be grouped by genus, from *Deltacoronavirus* to *Betacoronavirus* members (**Supp. Fig 1.A**). It is possible that the discovery of new *Ronaviridae* like viruses would help complement this emergent grouping by RNA genome size. The placement of AcNV in this context is intriguing; the similar and only other mollusk nido-like virus BNLV-1 was reported to have a much shorter 20.2 kb RNA genome. Nevertheless, there are some hints that may suggest that BNLV-1 could be longer than reported: The described BNLV-1 sequence NC_032496 does not present any stop codon interrupting the ongoing ORF1a, thus the suggested translation start site and the reported 5’UTR could correspond to a truncated coding region of ORF1aPP. In this direction, while nidoviruses ORF1a is typically much longer than ORF1b, ORF1a of BNLV-1 is about the same size as ORF1b. Perhaps future studies should revisit BNLV-1 genome sequence.

Given the estimated large size of AcNV genome, and the apparent constrains in terms of longer RNA viruses, the underlying basis of this discernible bar is under ongoing assessment. Probable causes of this size limit are under discussion, and the most plausible explanation could be associated with the high rate of RNA virus mutation associated with limited mechanisms of proof reading and repair, concomitant with high rates of replication (Holland, 1982; Denison et al., 2011). Supporting these assumptions, in parallel, there appears to be a correlation between small genome DNA viruses and high rate of mutation. In evolutionary terms, mutation rates could be associated not only to polymerase fidelity, but also to viral biology, genome architecture and to replication speed (Duffy et al., 2008; Campillo-Balderas et al., 2015). The size limit of RNA viruses is not specifically associated to monosegmented viruses. As a matter of fact, the largest members of the *Reoviridae* family, encompassing 9 to 12 dsRNA segments which added together grasp a total genome of ca. 32 kb, mirror the apparent size limit of *Nidovirales* and RNA viruses in general (Dolja & Koonin, 2018). From and evolutionary perspective, it was recently suggested that the nido-like clade of animal viruses emerged in early invertebrates and through horizontal virus transmission then colonized vertebrates. This proposition was based on the relative small genome size of deep-rooted arthropod nidoviruses, and the observation that the largest genomes correspond to vertebrate coronaviruses, postulating that gene incorporation and layering (and concomitant genome length growth) succeeded during the co-evolution of nidoviruses and vertebrates (Dolja & Koonin, 2018). The discovery here of the tentative AcNV associated to a mollusk, with the largest RNA genome of nidoviruses yet, with more affinity to intermediate-long *Roniviridae* than to larger coronaviruses, implies that the evolutionary history of *Nidovirales* could be more complex than expected.

### 3.3 A non-canonical genome organization of AcNV

One striking difference observed in AcNV and BNLV-1 genomes denotes a fundamental contrast to the predicted genome organization associated to nidoviruses, which consistently harbor at their 5’ region two large partially overlapping ORFs. Translation of ORF1a yields PP1a, and in a third of cases, codon slippage at the ORF1a/b overlap register the −1 reading frame, and continue with the translation of ORF1b (De Groot et al., 2012). On the other hand, BNLV-1 and AcNV appear to generate ORF1ab by means of suppression of termination of a premature stop codon after the ORF1a region. Perhaps it is worth mentioning here, that at a final mean coverage of 17,416X obtained of over 62 million viral RNA nt from 71 different *A. californica* studies, the proposed AcNV genome organization is highly supported and most probably not derived from a misassembly which could have resulted in the misidentification of overlapping ORFs and RFS signals. The identification of more divergent nido-like viruses may help to determine whether this is a conserved characteristic of mollusk associated nidoviruses or just anecdotal.

### 3.4 AcNV variability assessment should be complemented with additional data

Given the considerable number of viral reads, a tentative assessment of AcNV variability was generated. Polymorphic sites were defined with a conservative estimated frequency of at least ≥0.05 and a Phred score of 20 or higher to filter raw reads. Considering that sequencing errors of the diverse platforms reach at most ~1 % (Ion Torrent data), thus a 5% frequency threshold enables the detection of probable true variants in the virus RNA genome with high confidence. Cotten et al (2013), with a similar strategy, explored virus variants of human betacoronavirus using as threshold 1 %. A 5% frequency threshold was used by Van den Hoecke et al (2015) to assess the variation landscape of Influenza virus A. The detected variability among bioprojects appears to be non-significant or within the levels of intra-host variability. While dispersed in time, most NGS libraries were generated from lab reared animals from the same provider. Thus it is tempting to suggest (this is pure speculation) that the low intrinsic variability observed could be linked with a single virus lineage maintained inadvertently through the years at the breeder facilities.

### 3.5 AcNV has a broad tropism, is enriched in the CNS, and accumulates in neurons

The availability of a public collection of NGS RNA libraries of the model organism *A. californica*, allowed the tentative survey of the genomic RNA landscape of AcNV. Virus RNA transcripts were detected in an outstanding diversity of target tissue and organs, suggesting e pervasive tissue tropism, showing an emergent enrichment of AcNV in the CNS. Single-cell RNAseq data allowed the surveillance of AcNV at the cellular level, including not only specific types of neuron (e.g. abdominal ganglia neurons) but also the detection of AcNV in well-defined individual neurons of *A. californica* (e.g. R2, L29, R15). Interestingly, AcNV RNA accumulated at high levels in R2, which is the largest known neuron in the animal kingdom, offering an outstanding and unprecedented model for the study of virus dynamics at the single cell level. Besides their accessibility, ease to identify and size, each R2 neuron may yield over 1.9ug of RNA allowing to amplification-free RNA profiling at a single-neuron level (Moroz & Kohn, 2013). The standard of single-cell RNAseq preparation in any other organism cells, which have picogram levels of nucleic acids, is a robust amplification step, which may lead to irregular coverage, noise and imprecise quantification of sequencing data (Islam et al., 2011). The potential to conduct expression analyses on individual neurons of *A. californica* without the need of amplification, in naked-eye visible neurons, provide an elegant platform for genomics and physiological analysis within well-defined cellular networks: an ideal pathosystem to address single cell virus dynamics. It is tempting to suggest that the possibility of amalgamation of the study of this novel nido-like virus and a biological model system such as *A. californica*with a vast research toolkit, would allow the rapid deployment of modern molecular tools, which could redound in a rapid advancement in the biology of this entity. An impressive example of that is the recent integration of the house flies pathogen *Entomophtora muscae* into *Drosophila melanogaster*, which resulted in major advancements in the neurochemistry, genomics and neurobiological aspects of this fungal pathogen (Elya et al., 2017). In retrospective, it is perhaps worth contemplating the possibility that this hidden entity, present with certainty in samples of more than decade-old studies, might have influenced the outcome of any of the uncountable works published in the last 50 years. It is in this scenario that future studies should eagerly focus on the specific potential effects of this virus and its host. This invaluable information, missing in the literature, would shed light on the evolutionary history of the interaction of this virus and *A. californica*.

## 4. Materials and methods

### 4.1 Virus discovery

In order to assess the potential RNA virus landscape of *A. californica*, every available RNA resource at NCBI was employed for virus discovery. 117 NCBI SRA accessions of raw RNA sequencing reads were explored, obtained by 454 GS FLX and 20, Illumina HiSeq 2000, Illumina Genome Analyzer II and Ion Torrent PGM sequencing platforms, which are accessible at: https://www.ncbi.nlm.nih.gov/sra/?term=txid6500[Organism:noexp]. In addition, nine tissue-specific Transcriptome Shotgun Assemblies (TSA) were examined (GenBank accession numbers: GBCZ01000001, GBDA01000001, GBBV01000001, GBBW01000001, GBBE01000001, GAZL01000001, GBAV01000001, GBBG01000001, GBAQ01000001). Lastly, a collection of 255,605 *A. californica* expressed sequence tags (ESTs) of diverse developmental stages, tissues and treatments (Moroz et al, 2006; Fiedler et al., 2010) were retrieved from https://www.ncbi.nlm.nih.gov/nucest/?term=txid6500%5bOrganism:noexp]. The multiplatform RNA sequencing reads were de novo assembled with Trinity (version 2.5.1) https://github.com/trinityrnasea/trinityrnasea/wiki using standard parameters (Illumina data) and/or with the *de novo* assembly tool of CLC Genomics Workbench (v 8.0.0) with default settings (454 and Ion Torrent data). The obtained metatranscriptomes were assessed by bulk searches on a local server against a refseq virus protein database generated with data available at ftp://ftp.ncbi.nlm.nih.gov/refseq/release/viral/viral.1.protein.faa.gz. BLASTX with an expected value of 10e-5 was used as threshold, and hits were explored by hand. Tentative virus contigs were curated by iterative mapping of reads using Bowtie2 http://bowtie-bio.sourceforge.net/bowtie2/index.shtml virus Fragments Per Kilobase of transcript per Million mapped reads (FPKM) were estimated with Cufflinks 2.2.1 http://cole-trapnelllab.github.io/cufflinks/releases/v2.2.1/. The robust *A. californica* genome assembly AplCal3.0 (GenBank accession number GCF_000002075.1) was explored by TBLASTN searches (E-value >1e-1) using as query the predicted protein products of AcNV, in order to identify potential endogenous viral elements EVEs (Katzourakis & Gifford, 2010) with similarity with AcNV. In addition, raw DNA reads used to generate the *A. californica* genome (Bioproject PRJNA13635) were explored in detail by MEGABLAST to rule out potential unassembled EVEs missed in AplCal3.0 genome assembly.

### 4.2 Virus sequence annotation

Virus annotation was instrumented as described in Debat (2017), briefly, virus ORFs were predicted with ORFfinder (https://www.ncbi.nlm.nih.gov/orffinder/) translated gene products were assessed by InterPro (https://www.ebi.ac.uk/interpro/search/sequence-search) and NCBI Conserved domain database v3.16 (https://www.ncbi.nlm.nih.gov/Structure/cdd/wrpsb.cgi) to predict domain presence and architecture. Further, HHPred and HHBlits as implemented in (https://toolkit.tuebingen.mpg.de/#/tools/) were employed for domain prediction. Additionally, the Structural Classification of Proteins 2 tool available at (http://scop.mrc-lmb.cam.ac.uk/scop/) was used to complement annotation of divergent predicted proteins by hidden Markov models, and the Fold & Function Assignment server (http://ffas.sanfordburnham.org/ffas-cgi/cgi/document.pl) was used for the detection of remote homologies beyond the reach of other sequence comparison methods such as PSI-Blast. Pseudoknots were predicted with the Dotknot tool available at (http://dotknot.csse.uwa.edu.au/) and visualized with the VARNA 3.93 applet (http://varna.lri.fr/). Cleavage sites for signal peptides were predicted with the SignalP 4.1 Server (http://www.cbs.dtu.dk/services/SignalP). Prediction of transmembrane helices was performed with TMHMM v2.0 server (http://www.cbs.dtu.dk/services/TMHMM/). Intrinsic Protein Disorder regions were predicted with the disEMBL tool as implemented in http://dis.embl.de/ using standard parameters (predictors Remark-465, beta aggregation with Tango, low-complexity with CAST).

### 4.3 Phylogenetic insights

Phylogenetic insights based on predicted ORF1ab proteins were generated by MAFTT 7 https://mafft.cbrc.jp/alignment/software/ multiple amino acid alignments (BLOSUM62 scoring matrix) using as best-fit algorithm E-INS-I which drives local alignment with generalized affine gap costs, and thus it is applicable for large replicases where diverse domains are dispersed among several highly divergent regions. The aligned proteins were subsequently used as input for FastTree 2.1.5 maximum likelihood phylogenetic trees (best-fit model = JTT-Jones-Taylor-Thorton with single rate of evolution for each site = CAT) computing local support values with the Shimodaira-Hasegawa test (SH) and 1,000 tree resamples http://www.microbesonline.org/fasttree/. The FreeBayes v0.9.18S tool, using as minimum variant frequency ≥0.05 and additional standard parameters was employed for SNPs prediction https://github.com/ekg/freebayes. Results were integrated and visualized in the Geneious 8.1.9 platform (Biomatters Ltd.). Median haplotype networks were constructed with PopART version 1.7 (http://popart.otago.ac.nz) using the integer neighbor-joining algorithm based on MAFFT alignments of complete AcNV virus genome consensus sequences (>75% frequency call per-base) determined for *A. califonica* diverse Bioprojects.

### 4.4 Data availability

The AcNV virus (EK) RNA sequence determined in this study is available in GenBank under accession number MG878985 [not yet released to the public, is included as supplementary material of this manuscript submission.].

## Acknowledgements

I would like to express a sincere gratitude to the generators of the amazing underlying data used for this work: Dr. Leonid L. Moroz and Dr. Eric R Kandel. By following open access practices and supporting accessible raw sequence data in public repositories available to the research community, they and their teams have promoted the generation of new knowledge and ideas. Additional thanks to David Karlin for helpful comments. Special thanks to Elba Villanueva for critical reading of the manuscript.

## Funding

This research did not receive any specific grant from funding agencies in the public, commercial, or not-for-profit sectors.

## Conflicts of interest

The author declares no conflict of interest.

---

**Supplementary Figure 1.** Genome size in *Nidovirales* **A**) Intervals of RNA genome length in kb of families, sub-families, and genera of *Nidovirales* members. “Small nidoviruses” are represented by the *Arteviridae* family, “intermediate nidoviruses” by *Mesoniviridae*, and *Roniviridae* and *Coronaviridae* are considered “large nidoviruses”. **B**) Scaled genome diagrams of the 84 Refseq genomes of nidovirus available at NCBI and AcNV on top. Letters indicate taxonomic assignment to specific *Nidovirales* families or sub-families. The black line indicates the top 10 largest genomes, which are magnified in (**C**) AcNV EK sequence is over 2.4 kb longer than the largest ever reported RNA virus.

**Supplementary Figure 2.** Multinuclear coronavirus zinc-binding domain (CV-ZBD; Prosite =PS51653) ClustalW aa alignment of AcNV and similar nido-like viruses. While divergent, the predicted CV-ZBD domain of AcNV contains 10 His/Cys typical residues (asterisks) at equilocal conserved positions of several nidoviruses.

